# Cardiac hemodynamics and ventricular stiffness of sea-run cherry salmon (*Oncorhynchus masou masou*) differ critically from those of landlocked masu salmon

**DOI:** 10.1101/2022.04.06.487371

**Authors:** Yuu Usui, Misaki Kimoto, Akira Hanashima, Ken Hashimoto, Satoshi Mohri

## Abstract

Ventricular diastolic mechanical properties are important determinants of cardiac function and are optimized by changes in cardiac structure and physical properties. *Oncorhynchus masou masou* is an anadromous migratory fish of the Salmonidae family, and several ecological studies on it have been conducted; however, the cardiac functions of the fish are not well known. Therefore, we investigated ventricular diastolic function in landlocked (masu salmon) and sea-run (cherry salmon) types at 29–30 months post fertilization. Pulsed-wave Doppler echocardiography showed that the atrioventricular inflow waveforms of cherry salmon were biphasic with early diastolic filling and atrial contraction, whereas those of masu salmon were monophasic with atrial contraction. In addition, end-diastolic pressure–volume relationship analysis revealed that the dilatability per unit myocardial mass of the ventricle in cherry salmon was significantly suppressed compared to that in masu salmon, suggesting that the ventricle of the cherry salmon was relatively stiffer (relative ventricular stiffness index; *p* = 0.0263). Contrastingly, the extensibility of cardiomyocytes, characterized by the expression pattern of Connectin isoforms in their ventricles, was similar in both types. Histological analysis showed that the percentage of the collagen accumulation area in the compact layer of cherry salmon increased compared with that of the masu salmon, which may contribute to ventricle stiffness. Although the heart mass of cherry salmon was about 11-fold greater than that of masu salmon, there was no difference in the morphology of the isolated cardiomyocytes, suggesting that the heart of the cherry salmon grows by cell division of cardiomyocytes, but not cell hypertrophy. The cardiac physiological function of the fish varies with differences in their developmental processes and life history. Our multidimensional analysis of the *O. maosu* heart may provide a clue to the process by which the heart acquires a biphasic blood-filling pattern, i.e., a ventricular diastolic suction.

## Introduction

Ventricular stiffness, myocardial relaxation, and atrial pump functions are important for diastolic ventricular filling, a major determinant of cardiac output [1]. Ventricular stiffness is adjusted to suit the lifestyle of an individual and increases with age and growth [2, 3] by varying muscle mass, architecture, and geometry of the chambers [4]. Therefore, investigations and comparisons of the ventricular diastolic properties of individuals within a species at different developmental stages and body weights are crucial to advance the understanding of the physiological adaptations of the heart.

Ventricular diastolic functions at the organ and tissue level are evaluated by echocardiography and ventricular pressure–volume relationship analysis. For example, Doppler echocardiography shows blood inflow waveforms that determine the speed and direction of blood transfer from the atria to the ventricle using the pressure gradient between them [5]. Another is the assessment of mechanical properties for investigating the ventricular diastolic pressure–volume relationship, which has been used since early studies of cardiac mechanics [6–8]. End-diastolic pressure–volume relationship (EDPVR) analysis has been developed as one of the important concepts for interpreting cardiac mechanics and is established as a valid evaluation method for ventricular stiffness for both *in vivo* and *ex vivo* studies [4, 9–12]. EDPVRs on the graph are shown as a nonlinear curve, and the steeper the slope of this curve (dV/dP), the stiffer the ventricle. Clinical studies using EDPVR analytics have reported diastolic dysfunction with increasing ventricular stiffness by hypertension and aortic stenosis [13, 14]. When assessing the stiffness of ventricles of different sizes, their relative ventricular stiffness was also evaluated by normalization [15, 16]. Collagen accumulation, changes in collagen types, and inflammation were the distinguishing factors of stiffer ventricles [17–20].

Cardiomyocytes are heart-specific muscle cells and are responsible for generating diastolic and contractile force of the heart. The passive tension of cardiomyocytes is one of the determinants of ventricular filling dynamics as it provides resistance to expand the lumen of the ventricle. Connectin, encoded by *ttn*, is a protein comprising sarcomeres, and functions to generate tension forces in accordance with sarcomere lengths as biological springs [21–24]. The elastic potential of Connectin depends on the splicing of N2A and/or N2B spring segments, number of PEVK spring segment repeats, and phosphorylation levels of the protein [24–30]. Mammalian cardiomyocytes co-express the Connectin isoforms N2BA (3,700 kDa and compliant) and N2B (3,000 kDa and stiff) [31–34]. The expandability of cells changes according to the ratio of these isoforms [30, 35–37].

Teleosts possess a heart with four compartments: a sinus venosus, atrium, ventricle, and bulbus arteriosus (S1A and S1B Fig) [38–40]. Venous blood enters the sinus venosus from the liver and is pumped into the atrium and then the ventricle and is ejected to the ventral aorta through the bulbus arteriosus. Each compartment is separated by a valve. Most teleost ventricles are composed of two histologically distinct myocardial layers: a compact layer and a spongy layer [41–44]. The former is circumferentially arranged from dense cardiomyocytes as the ventricular wall under the epicardium, while the latter has cardiomyocytes arranged in a network, and is found in the luminal region of the ventricles. Venous blood from the atrium flows into the intertrabecular space. Based on the histological findings for compact layer thickness and the distribution patterns of the coronary vessels, fish ventricles are classified into four types (types I–IV) [44–46].

The Salmonidae family is extremely varied and includes 11 genera and at least 70 species that are believed to be extant [47]. Most species in this family change their habitats with migratory behaviors and have adapted to both freshwater and seawater environments. Owing to these ecological characteristics, their cardiovascular systems have been investigated as useful models for physiology [41, 48–52]. A triangular pyramidal ventricle is commonly found in the hearts of Salmonidae fish (Atlantic salmon [*Salmo salar*], sockeye salmon [*Oncorhynchus nerka*], and rainbow trout [*Oncorhynchus mykiss*]) [42, 53]. As the nutrient vessels at the ventricle are confined to the compact layer, these salmon ventricles were classified as type II. In addition, this ventricle type is believed to be an effective pump as it can transfer blood at higher heart rates than saccular and tubular ventricles [54, 55]. The salmonid compact layer thickens as it grows [56, 57]. Exercise, mechanical stress, and hormones also remodel the salmonid heart [58–64], and thermal changes induce collagen accumulation and increase heart stiffness [65, 66].

*Oncorhynchus masou masou* is an anadromous migratory fish belonging to the salmonid family and has two predominant life history types: landlocked (masu salmon) and sea-run (cherry salmon) (S1C Fig) [67, 68]. These fish have remarkably different skin patterns and body sizes. Transient increases in the secretion of hormones such as the thyroid hormone, growth hormone, cortisol, adrenocortical hormones, and sex hormones in the blood are involved in the smolt and anadromous processes [67–72]. While there have been many studies on the ecology of *O. masou*, to the best of our knowledge, there have been no studies to date on its cardiac functions.

In this study, we aimed to investigate the possible differences in the ventricular diastolic functions of masu salmon and cherry salmon. To compare the diastolic function of their hearts, we observed their atrioventricular inflows using pulsed-wave Doppler echocardiography and evaluated their passive ventricular mechanical properties by pressure–volume analysis *in vivo* and *ex vivo*; moreover, we compared their ventricular stiffness, heart histology, cardiomyocyte morphology, and the expression patterns of Connectin isoforms.

## Materials and methods

### Experimental approval

All experiments in this study were performed following the guidelines and approved protocols for animal care and use (approval number for animal use: 20-147) of the animal experiment committee at Kawasaki Medical School.

### Masu and cherry salmon

Twenty-one juvenile fish, six months post fertilization (mpf), were purchased online from Azuma Yougyojou Y.K. (Gunma, Japan). They were maintained for one week in 8-L freshwater tanks, with 10 fish per tank, because one juvenile died the day after transport. A total of 30 masu salmon (*O. masou* landlocked type) 29–30 mpf (25 fish) and 34 mpf (5 fish) were purchased online from Utanogawa Yamame Yougyojou (Yamaguchi, Japan). After arrival, they were maintained in 70-L freshwater tanks, with 5 or 10 fish per tank. They were used in the experiment within one week of their arrival. Eleven cherry salmon (*O. masou* sea-run type) at 29–30 mpf were purchased from Wakao-Suisan Co., Ltd. (Hyogo, Japan). After transport, they were maintained in 200-L seawater tanks with a salinity of 35 g/L (1.023 specific gravity) [73], and two or three fish per tank. For the cherry salmon, *in vivo* experiments were performed within two days of their transport. All fish tanks were maintained at 10°C with a 14-/10-h light/dark cycle. The salmon were fed a commercial diet (Remix, Meito Suien Co., Ltd., Aichi, Japan, Cat# M-450) and promptly euthanized after data collection. Fish were randomly selected for this study as sex could not be specified at the time of purchase.

### Echocardiography and electrocardiography

Echocardiography in pulsed-wave Doppler mode and electrocardiography were recorded using an Aplio 300 system (Toshiba Medical System Corp., Tochigi Japan) with a 14-MHz transducer and three electrodes. Mild anesthetic induction by MS-222 (50 mg/L, Sigma-Aldrich, Merck KGaA, Darmstadt, Germany) [74] was used to treat five masu salmon and two cherry salmon until there was minimal gill movement. The anesthetized fish were placed upside down in the tank. The ultrasound transducer probe was placed on their heart positions, and the detection depth and brightness were adjusted for each fish (S2A Fig). The waveforms of atrioventricular inflows were observed immediately after ventricular outflow recordings because it is impossible to measure these flows simultaneously with a single probe. From the pulse-wave Doppler images, ventricular outflow times and velocities were calculated using Fiji software version 2.3.0 [75]. Each interval between the end of the ventricular outflow and the onset of the first atrioventricular inflow was calculated by subtracting the average time of ventricular outflow from each time from the peak of the R wave to the onset of atrioventricular inflow. The changes in the potential across the *O. masou* body surface by electrical excitation of the heart were measured by clipping electrodes to their pectoral and pelvic fins (S2A Fig). Aeration was performed during the measurements. After recording, all fish were euthanized using MS-222 (250 mg/L) for organ sampling.

### *In vivo* atrial and ventricular pressure measurements

Five anesthetized masu salmon 34 mpf were placed upside down in the tank. A Single Transducer Set, DTX Plus DT-4812 (Argon Medical Devices, Inc., Plano, TX, USA), with needles was used to measure the pressure. Using an echocardiographic live image, a 25-G needle was inserted from the ventral side into the ventricle and a 27-G needle was inserted under the operculum into the atrium. The ventricular pressure tracing data were used to mediate the pressure recording interface (Power Lab 8/30, ADInstruments Pty Ltd., Dunedin, New Zealand) and were digitized at 200 Hz using LabChart 7 software (ADInstruments) and recorded. After recording, all fish were euthanized using MS-222 (250 mg/L).

### *Ex vivo* ventricular pressure–volume analysis

Four masu salmon and three cherry salmon were euthanized, after which their hearts were removed. A catheter was inserted into the ventricles through the atrioventricular valve and fixed by ligation. Fluids in the ventricular lumen were washed using 10 U/mL heparin FUJIFILM Wako Pure Chemical Corp., Osaka, Japan) and 10 mM 2,3-butanedione monoxime (BDM; FUJIFILM Wako Pure Chemical Corp.) in Ca^2+^- and Mg^2+^-free phosphate-buffered saline (PBS; Takara Bio Inc., Kusatsu, Japan). The aortic valve was then ligated. To measure the pressure–volume relationship of the ventricles, saline solution was injected into the ventricles using an infusion pump (Terumo Corp., Tokyo, Japan). The ventricular pressure data were recorded at 400 Hz with a multielectrode conductance catheter using LabChart 7 software The data for each EDPVR were described as an exponential fit based on the following equation:

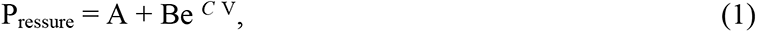

where A, B, and *C* were constants that describe the ventricular exponential pressure-normalized volume property; constant A indicated the intercept on the pressure axis (A = -B), and *C* indicated the index of the ventricular stiffness. V was the ventricular volume [4]. When comparing the ventricular stiffness in hearts of different sizes, V was normalized by the ventricular mass to calculate the stiffness per unit of ventricular mass and described by the following equation:

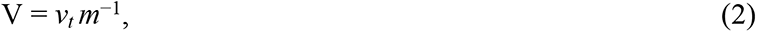

where *v_t_* was the total volume of saline infused into the ventricle at time *t*, *v*_0_ = 0, and *m* was the ventricular mass [15].

### Tissue staining

Masu and cherry salmon hearts were fixed in 4% paraformaldehyde (PFA; Sigma-Aldrich, Merck) at 4°C for 48 h and embedded in paraffin blocks. The mounted glass slides with 5-µm-thick heart sections were immersed in Clear Plus (FALMA Corp., Tokyo, Japan) followed by 100%, 95%, and 70% ethanol solutions for deparaffinization, and finally rinsed with water. The methods for Elastica van Gieson and hematoxylin and eosin staining are described in the sub-sections. After staining, the tissue sections were dehydrated in 70%, 95%, and 100% ethanol solutions and ethanol/xylene mixed solution, immersed in xylene four times, and sealed using a mounting medium (Muto Pure Chemicals Co., Ltd., Tokyo, Japan). The sections were observed under a bright-field microscope IX53, (Olympus, Tokyo, Japan) using the cellSens software version 1.18 (Olympus) and BZ-X700 microscope (Keyence, Osaka, Japan). The collagen fiber content in the compact layer and cardiomyocyte density were measured using Fiji software version 2.3.0 [75].

### Elastica van Gieson staining

The heart sections were immersed in Maeda’s resorcinol fuchsin stain solution (Muto Pure Chemicals Co., Ltd.) for 1 h and washed with 100% ethanol. The sections were then immersed in Weigert’s iron hematoxylin solution and washed with Milli-Q water. The specimens were then immersed in Picro–Sirius Red solution (ScyTek Laboratories, Inc., West Logan, UT, USA) for 15 min, air-dried, and rinsed with xylene.

### Hematoxylin and eosin staining

The heart sections were immersed in Hansen’s hematoxylin solution (Sigma-Aldrich, Merck) for 5 min, followed by washing with water. To define the nuclei, sections were immersed in 0.2% hydrochloric acid–70% ethanol solution for 1 min, and then immersed in eosin Y solution (Millipore, Merck) for 30 s.

### Cardiomyocyte isolation and primary cultures

For the isolation and primary cultures of the cardiomyocytes from the masu and cherry salmon, we used a previously described zebrafish protocol [76]. We euthanized two masu salmon 29– 30 mpf and two cherry salmon 29–30 mpf, collected their ventricles, and washed their lumens by injecting heparin buffer (10 U/mL heparin in PBS). The ventricles were dissected into 3-mm^2^ columns and shaken at 800 rpm for 90–120 min at 30°C with 750 µL digestion buffer (12.5 µM CaCl_2_, 5 mg/mL collagenase type II [Thermo Fisher Scientific Inc., Waltham, MA, USA], 5 mg/mL collagenase type IV [Thermo Fisher Scientific Inc.], 10 mM HEPES, 30 mM taurine, 5.5 mM glucose, and 10 mM BDM). Next, the cell suspension was filtered through a 100-µm nylon cell strainer, and 1 mL CaCl_2_ buffer (12.5 µM CaCl_2_, 10% fetal bovine serum [FBS; HyClone, Cytiva, Marlborough, MA, USA, Lot#15N353], 10 mM HEPES, 30 mM taurine, 5.5 mM glucose, and 10 mM BDM) was added and centrifuged at 200 × *g* for 5 min at 4°C. After discarding the supernatant, 1 mL of CaCl_2_ buffer was added and the mixture was centrifuged at 200 × *g* for 5 min at 4°C. After being centrifuged, the isolated cells were resuspended using the culture media (Eagle’s minimum essential medium with 4.5 g/L glucose, 2 mM L-glutamine, 1 mM sodium pyruvate, and phenol red [FUJIFILM Wako Pure Chemical Corp.], 5 mM BDM, 5% FBS, and 100 U/mL penicillin-streptomycin [Thermo Fisher Scientific Inc.]) and seeded on a glass-bottom dish (Iwaki Co. Ltd., Tokyo, Japan) with 0.01% poly-L-lysine (Millipore, Merck) coating. Primary cells were cultured at 25°C under 5% CO_2_. Twenty-four hours after incubation, the primary cells were observed using a BZ-X700 fluorescence microscope (Keyence). The longitudinal length and area of the cardiomyocytes were measured using Fiji software version 2.3.0 [75].

### Immunostaining

Primary cardiomyocytes were fixed in 4% PFA at 4°C for 15 min. They were then permeabilized with 0.02% Triton X-100 in PBS for 15 min and blocked with 5% bovine serum albumin (Sigma-Aldrich, Merck) in PBS for 3 h. These cells were incubated with primary antibody, monoclonal anti-α-actinin (Sarcomeric) antibody produced in mouse (1:1000, Sigma-Aldrich, Merck, clone EA-53, Cat# A7811), at 4°C overnight, and subsequently stained with a secondary antibody, polyclonal goat anti-mouse IgG (H+L) cross-adsorbed secondary antibody, Alexa Fluor 488 (1:1000, Thermo Fisher Scientific Inc., Cat# A-11001), and Hoechst33342 (Tokyo Chemical Industry Co., Ltd., Tokyo, Japan, Cat# H342) for 1 h at 25°C. Using an FV1000 confocal laser scanning microscope system mounted IX81 (Olympus), the sarcomeric structures and the number of nuclei in the cardiomyocytes were then examined.

### Sodium dodecyl sulfate-agarose gel electrophoresis and western blotting

The atria, ventricles, and bulbus arteriosus were separated from the hearts of the masu and cherry salmon 29–30 mpf. Each sample was homogenized using Polytron (Kinematica AG, Malters, Switzerland) in a sample buffer (8 M urea, 2 M thiourea, 3% sodium dodecyl sulfate [SDS], 75 mM dithiothreitol, 0.03% bromophenol blue, 0.05 M Tris-HCl [pH 6.8], and protease inhibitor cocktail [Thermo Fisher Scientific Inc.]), heated at 65°C for 10 min, centrifuged at 16,000 × *g* for 5 min at 4°C and then the supernatant was collected. Cleared lysates were loaded into wells of the SDS-agarose gels (1% SeaKem Gold Agarose [Lonza, Basel, Switzerland], 30% glycerol, and 1 × Tris/glycine/SDS [TG-SDS] buffer [Takara Bio Inc.]) and then electrophoresed in a running buffer (1 × TG-SDS buffer and 10 mM 2-mercaptoethanol) at 0.01 A for 90 min [77]. To avoid the gel sliding from the gel plate during manipulation, the SDS-agarose gel was stacked and fixed on a 1-cm-high acrylamide plug gel (12% acrylamide, 10% glycerol, 0.5 M Tris-HCl [pH 9.0], 0.0015% N,N,N′,N′-tetramethylethylenediamine, and 0.056% ammonium persulfate). The patterns of the band peaks on the Coomassie Brilliant Blue (CBB)-stained gels were scanned using the Gel Analyzer plugin of Fiji software version 2.3.0.

For western blotting, the electrophoresed proteins were transferred from SDS-agarose gel to a nitrocellulose membrane (Bio-Rad Laboratories, Inc., Hercules, CA, USA) with a semi-dry western blot system (Pierce Power Blotter, Thermo Fisher Scientific) at 25 V for 10 min. After blocking the membrane with Tris buffer saline containing 0.05% Tween 20 and 5% skim milk, the Connectin proteins in the heart tissue were identified using a polyclonal primary antibody, the C-terminus of chicken Connectin (1:1000, Pc72C) [78], and polyclonal goat anti-rabbit immunoglobulin/HRP (1:1000, DAKO Agilent Technologies, Inc., Palo Alto, CA, USA, Cat# P044801-2) as a secondary antibody. Connectin blots were detected using the chemiluminescence reagent Western Lightning ECL Pro (PerkinElmer Co., Ltd., Waltham, MA, USA) and acquired using ImageQuant LAS 4000 (GE Healthcare, Chicago, IL, USA).

### Transmission electron microscopy

To observe the sarcomere structures in the cardiomyocytes using a transmission electron microscope (TEM), the fish hearts at 29–30 mpf were washed with 10 U/mL heparin and 10 mM BDM in PBS and fixed with 4% PFA and 2.5% glutaraldehyde in PBS for 1 h at 25°C. The fixatives were thoroughly washed with PBS. The samples were then post-fixed in 1% osmium tetroxide in PBS and immersed in 50%, 60%, 70%, 80%, 90%, 95%, 99%, and 100% ethanol consecutively for dehydration. Next, the samples were passed through propylene oxide, embedded in epoxy resin, and polymerized in an incubator for 72 h at 60°C. Sections that were 500 nm thick were produced from the resin block using an ultramicrotome and collected on glass slides. After reaching the desired section, 70-nm ultrathin sections were cut and placed on a single-hole grid coated with Formvar film. The grids were air-dried and stained with uranyl acetate and lead citrate. Electron micrographs were recorded using TEM JEM-1400 (JEOL Ltd., Tokyo, Japan), operated at 80 kV.

### Statistical analysis

The experimental results are presented as the mean ± standard deviation (SD); the number of samples used in each analysis is indicated in each figure legend. *P*-values were calculated using Student’s *t*-test or one-way analysis of variance (ANOVA) with Tukey’s multiple comparisons test and are indicated on graphs and text. The threshold for significant difference was set at a *p*-value of ≤ 0.05. GraphPad Prism8 software version 8.4.3 (GraphPad Software, San Diego, CA, USA) was used to conduct all statistical analyses.

## Results

### Evaluation of the ventricular hemodynamics in masu and cherry salmon with pulsed-wave Doppler echocardiography

At 29–30 mpf, the cherry salmon were found to be approximately 2.4 times bigger and 17 times heavier than the masu salmon (Fig 1A and 1B; Table 1). We also investigated their hemodynamics at this developmental stage. Pulsed-wave Doppler echocardiography and electrocardiography were simultaneously performed to observe their atrioventricular inflow in sinus rhythms. To set the pulsed-wave Doppler gate window downstream of the atrioventricular valve, a sagittal long-axis view including the atrium, ventricle, and bulbus arteriosus was acquired (S2B and S2C Fig; S1 and S2 Movies). Electrocardiography revealed regular P waves, QRS complexes, and T waves (S2D Fig).

**Fig 1.**
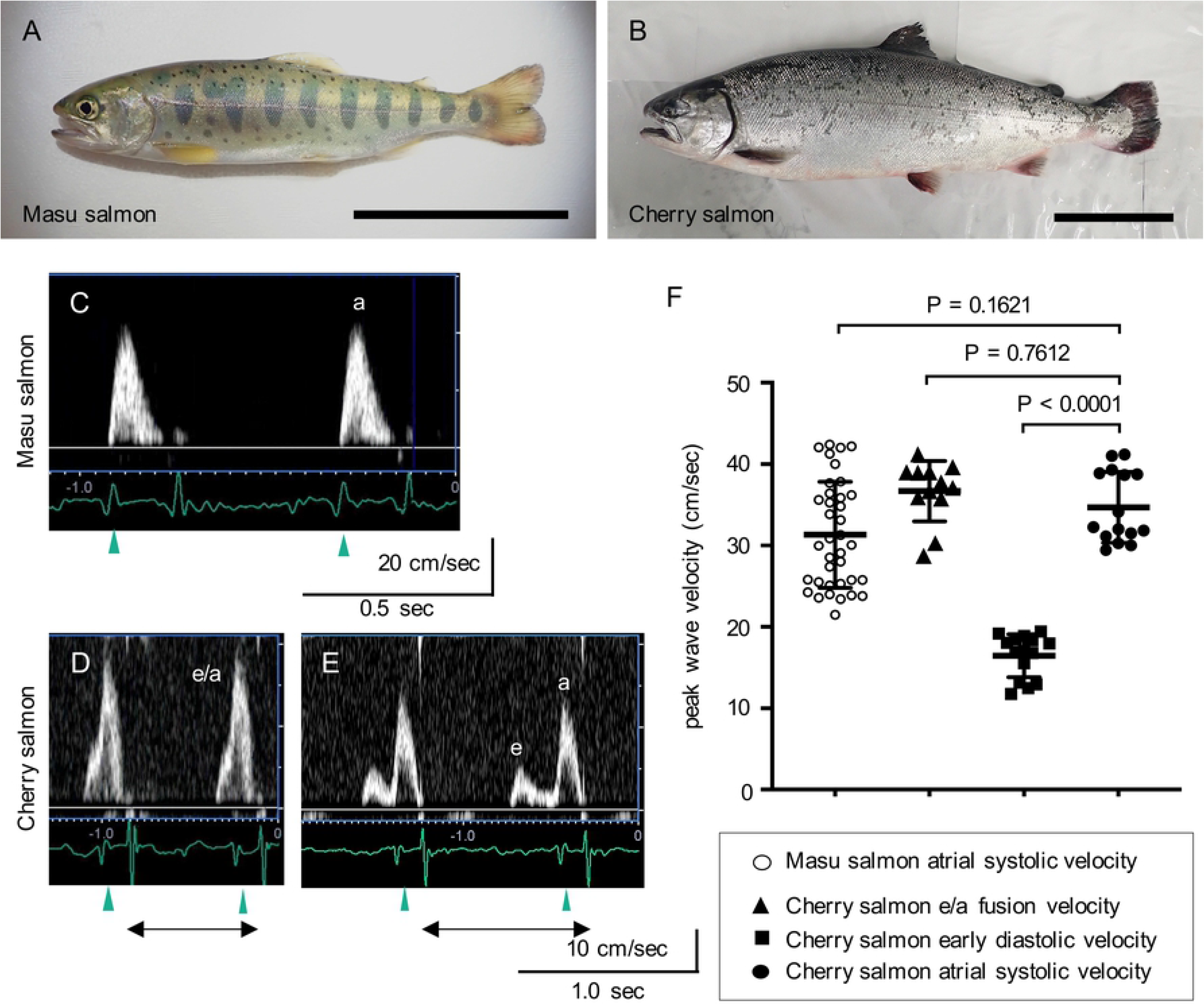
Recordings of the waveforms of the atrioventricular inflows in masu and cherry salmon using pulsed-wave echocardiography. (A, B) Representative *Oncorhynchus masou masou* pictures: (A) masu salmon at 29 months post fertilization, (B) cherry salmon reared in seawater for five months after being reared in freshwater for 24 months. Scale bars = 10 cm. (C, D, E) Representative velocity waveforms of atrioventricular inflows in the masu (C) and cherry salmon of monophasic (D) and biphasic patterns (E). Blood passing the atrioventricular valve was observed using pulsed-wave Doppler echocardiography simultaneously with electrocardiography. Green arrowheads indicate P waves. Black left–right double arrows indicate R-R intervals. a: atrial systolic velocity, e/a: fusion velocity of atrial systolic and early diastolic inflows, e: early diastolic velocity. (F) Peak wave velocity. ○: masu salmon atrial systolic velocity (31.34 ± 6.43 cm/s, N = 37 peaks from five fish), ▲: cherry salmon early diastolic waves and atrial systolic waves (e/a) fusion velocity (36.69 ± 3.57 cm/s, N = 12 peaks from two fish), ▪: cherry salmon early diastolic velocity (16.47 ± 2.56 cm/s, N = 15 peaks from two fish), ●: cherry salmon atrial systolic velocity (34.70 ± 4.22 cm/s, N = 15 peaks from two fish). Lines and error bars indicate the mean ± standard deviation.

**Table 1.**
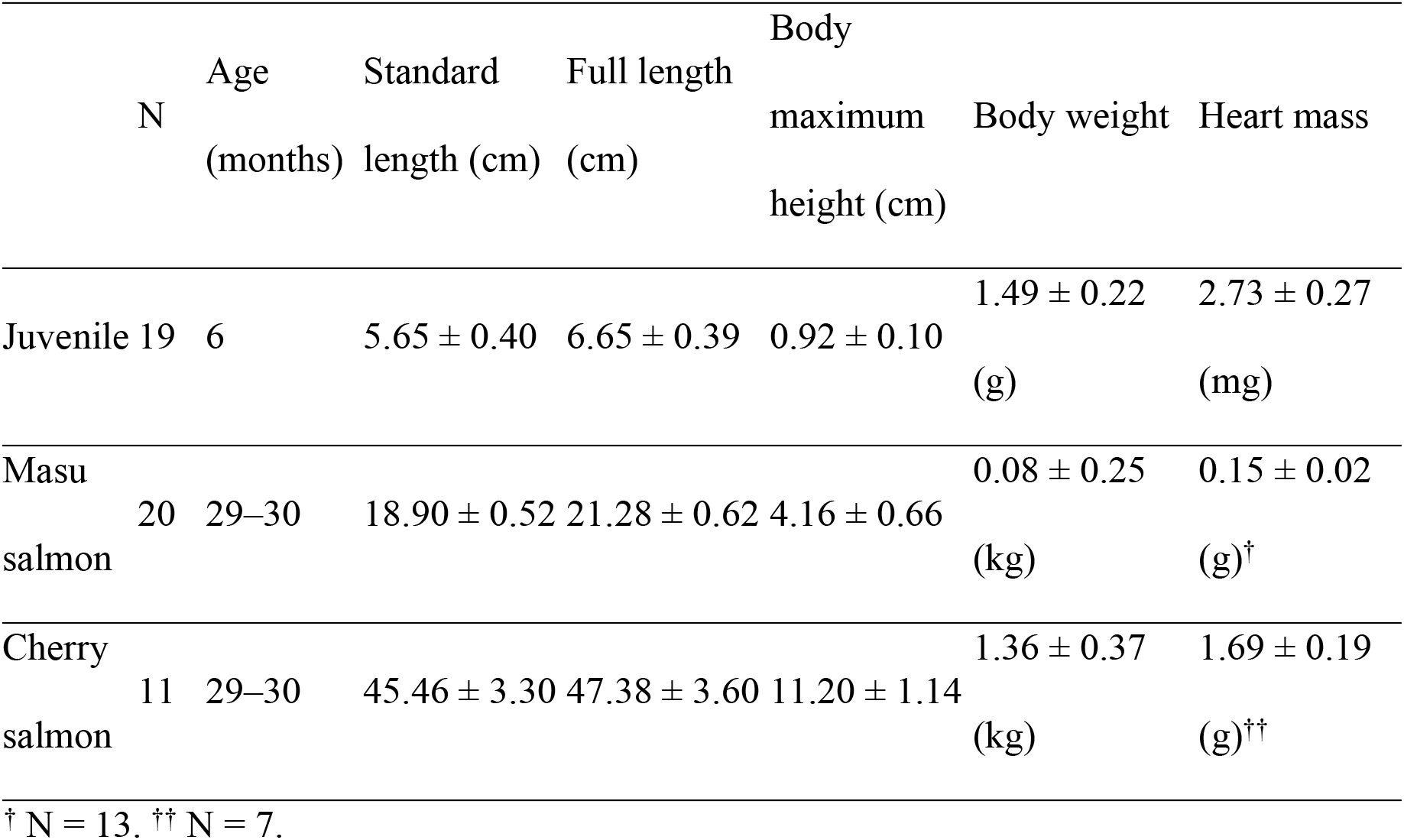
Body size and heart mass measurements (means ± standard deviation) for *Oncorhynchus masou masou*.

Pulsed-wave Doppler echocardiography revealed that the atrioventricular inflow patterns in the masu salmon showed sequential single forward flow waveforms that were synchronized with the P waves, indicating atrial contraction (Fig 1C). The heart rate of masu salmon as recorded with echocardiography was 109 ± 12.0 beats/min (bpm, N = 5 fish) (S1 Table). In masu salmon #5, the time from the R wave to the end of ventricular outflow was 326.75 ± 1.04 ms (N = 4 beats) (S3A Fig). When the interval times for the onset of ventricular systole (R-R interval times) took 624 ms (short) and 958 ms (long), the time between the end of ventricular outflow and the appearance of atrioventricular inflow waveforms was estimated to be 160 ms and 396 ms, respectively (S3 Fig). Regardless of the duration of the R-R interval, the atrioventricular inflow waveforms showed a monophasic pattern; furthermore, the atrial pressure in the ventricular diastole of the masu salmon was the same as the ventricular pressure for a while after isovolumic relaxation and became higher than the ventricular pressure in the atrial systole (S4 Fig). Hence, the monophasic inflow waveforms in masu salmon were presumed to indicate atrial systolic velocity.

Interestingly, the forward flow waveforms began to appear in the atrioventricular inflow patterns of the cherry salmon, even before the P waves were recorded (Fig 1D). When the R-R interval took > 0.77 s, echography recorded clear biphasic waveforms (Fig 1E, S3 Fig). The low forward waveforms appeared before the P wave, and the high forward waveforms appeared after the P wave. Therefore, the former and the latter were presumed to be comparable to the early ventricular diastolic and atrial systolic velocities, respectively. In addition, the sequential monophasic waves in cherry salmon were likely to be the fusion of these two velocities (early ventricular diastolic and atrial systolic [e/a] fusion velocity). The mean ± SD of the early diastolic velocity peak in the cherry salmon was 16.47 ± 2.56 cm/s, which was lower than the atrial systolic velocity (*p* < 0.0001, Fig 1F). The ratio of the early diastolic velocity/atrial systolic velocity (E/A ratio) was 0.48 ± 0.06 in the cherry salmon. In cherry salmon #1, the interval time from the R wave to the end of ventricular outflow was 351.75 ± 1.09 ms (N = 4 beats). Whether the R-R interval times were 720 ms (short), 772 ms (intermediate), or 908 ms (long), the times from the end of ventricular outflow until the appearance of atrioventricular inflow waveforms remained almost the same, at 92–96 ms (S3 Fig). Under MS-222 anesthesia, the heart rates of cherry salmon were 72 ± 5.3 bpm (N = 2 fish) and lower than that of masu salmon (*p* = 0.0164, S1 Table).

### Analysis of ventricular stiffness in masu and cherry salmon

To investigate ventricular stiffness, we performed the EDPVR analysis *ex vivo*. The interventricular pressure of masu salmon reached 20 mmHg in response to injection with 0.27– 0.31 mL saline (S4A Fig), while that of cherry salmon reached ∼20 mmHg in response to injection with 1.13–1.91 mL saline, and the smaller ventricles tended to show steeper EDPVR curves (S4 Fig). However, the normalized EDPVR curves of the cherry salmon shifted to the left were steeper compared to those of the masu salmon (Fig 2A). The indices of ventricular stiffness were 1.042 ± 0.187 in masu salmon and 2.205 ± 0.736 in cherry salmon, and the ventricles of cherry salmon were significantly stiffer than those of masu salmon based on the measurement of ventricular stiffness per unit myocardial mass (*p* = 0.0263, Fig 2B).

**Fig 2.**
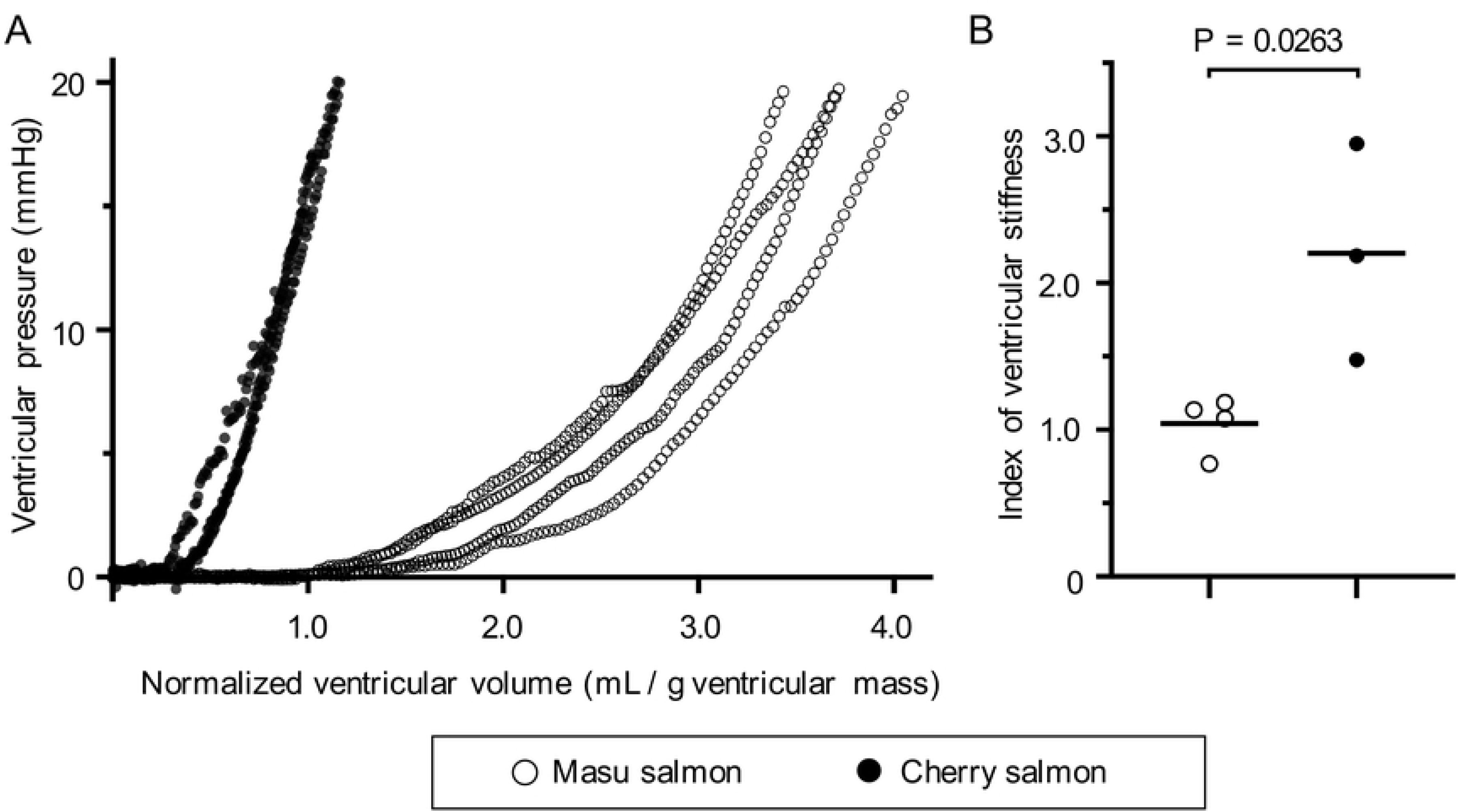
Relative ventricular stiffness of masu and cherry salmon. (A) The normalized end-diastolic pressure–volume relationship (EDPVR) at 30 months post fertilization for masu and cherry salmon. The horizontal axis shows the ventricular volume normalized by the mass of the masu (N = 4) and cherry salmon ventricles (N = 3). The vertical axis shows the ventricular pressure. Ventricular mass is shown in S5B Fig. (B) Scores for ventricular stiffness in the masu and cherry salmon. The results of the EDPVR (A) were appended in Equation (1) to obtain the exponential *C*, and the means ± standard deviations were calculated; masu salmon (*C* = 1.042 ± 0.187), cherry salmon (*C* = 2.205 ± 0.736). Black horizontal lines in (B) indicate the means. ○: masu salmon, ●: cherry salmon.

### Morphological and histological analysis of the hearts from masu and cherry salmon

To investigate the cause underlying the different ventricular diastolic properties between the masu and cherry salmon, hearts from these fishes were morphologically and histologically examined. At 29–30 mpf, the ventricles of both fish displayed a pyramidal shape (Fig 3A and 3B). The heart mass of cherry salmon was approximately 11 times greater than that of masu salmon (Table 1). The juvenile ventricles at 6 mpf also exhibited pyramidal shapes (S1D and S1E Fig). We then observed the Elastica van Gieson-stained and hematoxylin and eosin-stained sections of masu and cherry salmon hearts (Fig 3C–J and S6 Fig). Their ventricles had at least two layers of myocardium, i.e., a compact layer and a spongy layer (Fig 3C, 3D, 3G, and 3H). The compact layer in the myocardium of cherry salmon had more coronary vessels compared with that in masu salmon (Fig 3D and 3H). Further, this layer was thicker in cherry salmon than that in masu salmon (Fig 3D and 3H), and the ratios of the compact layer area to the total ventricular area on sagittal heart slices were significantly different between masu (28.95% ± 5.14%) and cherry salmon (39.56% ± 2.50%; *p* = 0.0387, Fig 3K). The cell density in the compact layer was 6297.3 ± 762.4 and 6733.7 ± 1020.7 nuclei/mm^2^ in masu and cherry salmon, respectively, and there was no significant difference between masu and cherry salmon (*p* = 0.3177, Fig 3L).

**Fig 3.**
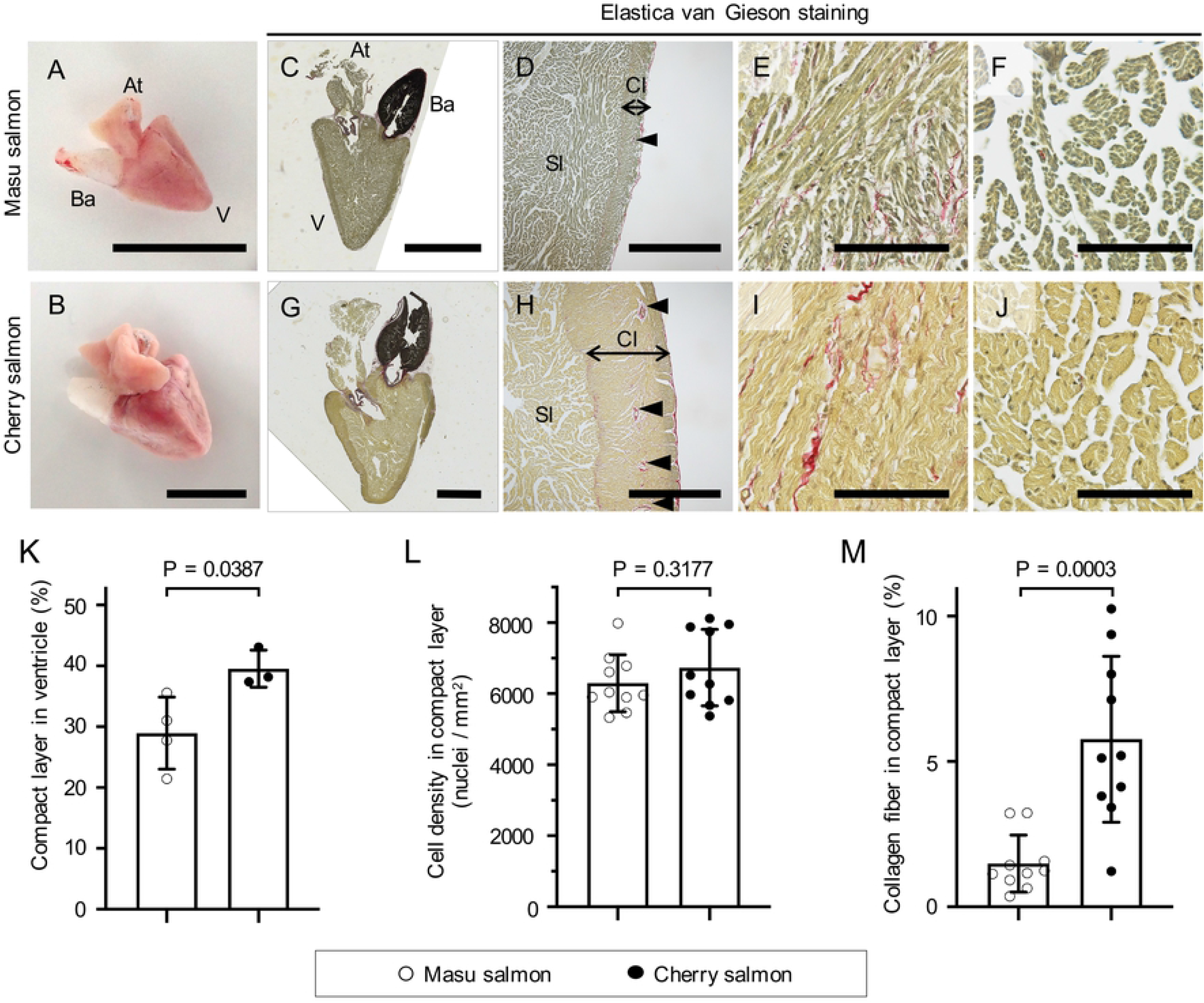
Histological analysis of masu and cherry salmon hearts. (A, B) Representative images of masu (A) and cherry (B) salmon hearts 29 months post fertilization (mpf). At: atrium, V: ventricle, Ba: bulbus arteriosus. (C–J) Elastica van Gieson staining images of the sagittal sections of the hearts of masu and cherry salmon 29 mpf. (C, G) Images of the whole hearts. (D, H) Magnified images of the ventricles. Black double arrows indicate the thickness of the compact layer. Black arrowheads indicate coronary vessels. (E, I) Higher magnification images of the compact layers. (F, J) Higher magnification images of the spongy layers. Collagen fibers: bright red; cytoplasm: yellow; elastic fibers: purple-black; nuclei: dark black. Cl: compact layer, Sl: spongy layer. Scale bars = 1 cm in (A) and (B); 5 mm in (C) and (G); 1 mm in (D) and (H); 100 µm in (E), (I), (F), and (J). (K) Percentage compact layer in the sagittal ventricle sections; masu salmon (28.95% ± 5.14%, N = 4), cherry salmon (39.56% ± 2.50%, N = 3). (L) The number of nuclei per yellow area in the compact layer; masu salmon (6297.3 ± 762.4 nuclei/mm^2^, N = 10), cherry salmon (6733.7 ± 1020.7 nuclei/mm^2^, N = 10). (M) The collagen fiber area percentage in the compact layer; masu salmon (1.492% ± 0.932%, N = 10), cherry salmon (5.768% ± 2.708%, N = 10). Bar graphs and error bars indicate the means and standard deviations, respectively. ○: masu salmon, ●: cherry salmon.

The extracellular matrix is one of the parameters used to adjust tissue compliance. For example, elastin is an extracellular matrix that provides tissues with high levels of elasticity [79]. Bulbus arteriosus is an elastin-rich tissue [80] and stains purple–black with Elastica van Gieson staining (Fig 3C and 3G). The collagen fibers were tinted an intense red with Elastica van Gieson staining. Investigation of the heart sections of masu and cherry salmon revealed that the compact and spongy layers included collagen fibers (Fig 3E, 3F, 3I and 3J). The epicardium, which is the outermost layer of the heart, the area around the coronary vessels, and the contact areas of the compact and spongy layers were found to be rich in collagen fibers (Fig 3D and 3H). The collagen fibers occupied 1.492% ± 0.932% of the compact layer in the ventricles from masu salmon and 5.768% ± 2.708% of the compact layer in the ventricles from cherry salmon, and the latter exhibited a significant accumulation of collagen fibers (*p* = 0.0003, Fig 3M).

Hypertrophy of the ventricular wall in response to hypertrophic cardiomyopathy is one of the factors that impairs the diastolic function of ventricles [81, 82]. Importantly, hypertrophy of the ventricular tissue is sometimes accompanied by an increase in cardiomyocyte size [83]. Mammalian cardiomyocytes become larger and maturate via multinucleation and polyploidization after birth [84, 85]. To investigate whether cell hypertrophy is involved in heart growth, the cardiomyocytes of masu and cherry salmon were isolated, and their morphologies were analyzed. Most of the isolated cells that were attached to the dish showed spindle or rectangular shapes (Fig 4A and 4B). These cells were stained in a band pattern using the cardiomyocyte marker α-actinin 2 (S7A–D Fig). Smaller and round-shaped cells were negative for α-actinin 2 (S7E and S7F Fig). The longitudinal length and area of the cultured primary cardiomyocytes were not significantly different between masu (56.30 ± 18.78 µm and 559.07 ± 238.92 µm^2^) and cherry salmon (57.63 ± 19.75 µm and 518.41 ± 197.03 µm^2^; *p* = 0.7259 and *p* = 0.3510; Fig 4C and 4D, respectively). In these primary culture experiments, we found one binucleated cardiomyocyte in cherry salmon, and the rest of the cardiomyocytes were mononucleated (S8 Fig).

**Fig 4.**
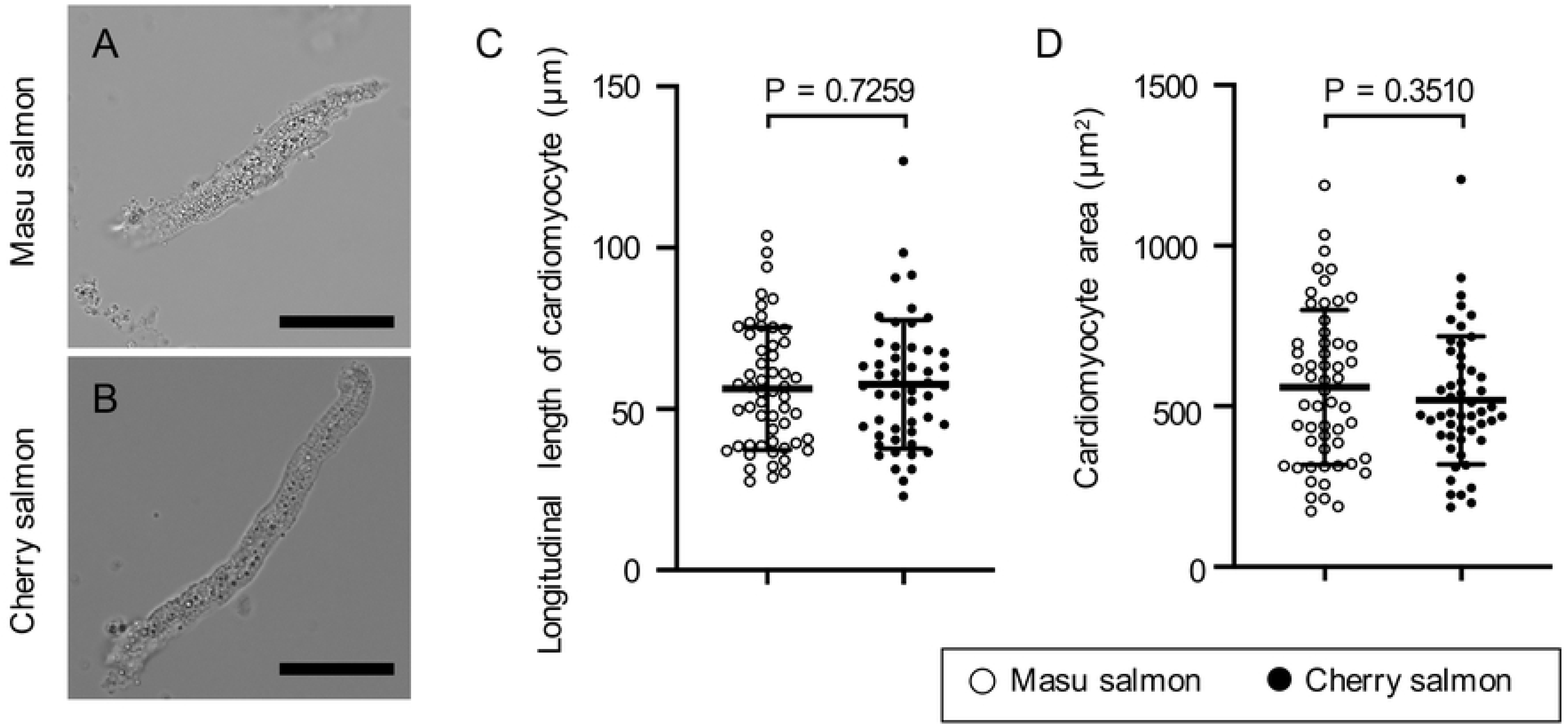
Morphology and sizes of the isolated cardiomyocytes. (A, B) Representative bright-field images of the primary cultured cardiomyocytes of masu and cherry salmon 29 months post fertilization. Scale bars = 20 µm. (C) Length of the long axis of the isolated cardiomyocytes; masu salmon (56.30 ± 18.78 µm, N = 55), cherry salmon (57.63 ± 19.75 µm, N = 50). (D) Area of the isolated cardiomyocytes; masu salmon (559.07 ± 238.92 µm^2^, N = 55), cherry salmon (518.41 ± 197.03 µm^2^, N = 50). Lines and error bars indicate the mean ± standard deviation. ○: masu salmon, ●: cherry salmon.

### Connectin expression patterns in the hearts of masu and cherry salmon

The expression patterns of connectin isoforms affect ventricular diastolic function [30, 35–37]; these patterns are regulated by thyroid hormone in mammals [86]. In *O. masou*, increased secretion of this hormone induces downstream migratory behavior and smoltification [69, 72]. From these assumptions, we hypothesized that masu and cherry salmon exhibit different expression patterns for cardiac Connectin isoforms. To investigate the cardiac Connectin expression patterns, SDS-agarose gel electrophoresis was performed (Fig 5A). Band pattern analysis of the CBB-stained gel revealed the presence of two peaks in the α-connectin (T1) zone, indicating expression of intact Connectin in the ventricles of both masu and cherry salmon (Fig 5B); furthermore, western blotting with the anti-connectin antibody [78] confirmed that these bands corresponded to the connectin from masu and cherry salmon (S9 Fig). These results suggest that *O. masou* at 29–30 mpf expressed Connectin (∼3,700 kDa) in their ventricles and that there was no difference in the expression pattern of Connectin between masu and cherry salmon.

**Fig 5.**
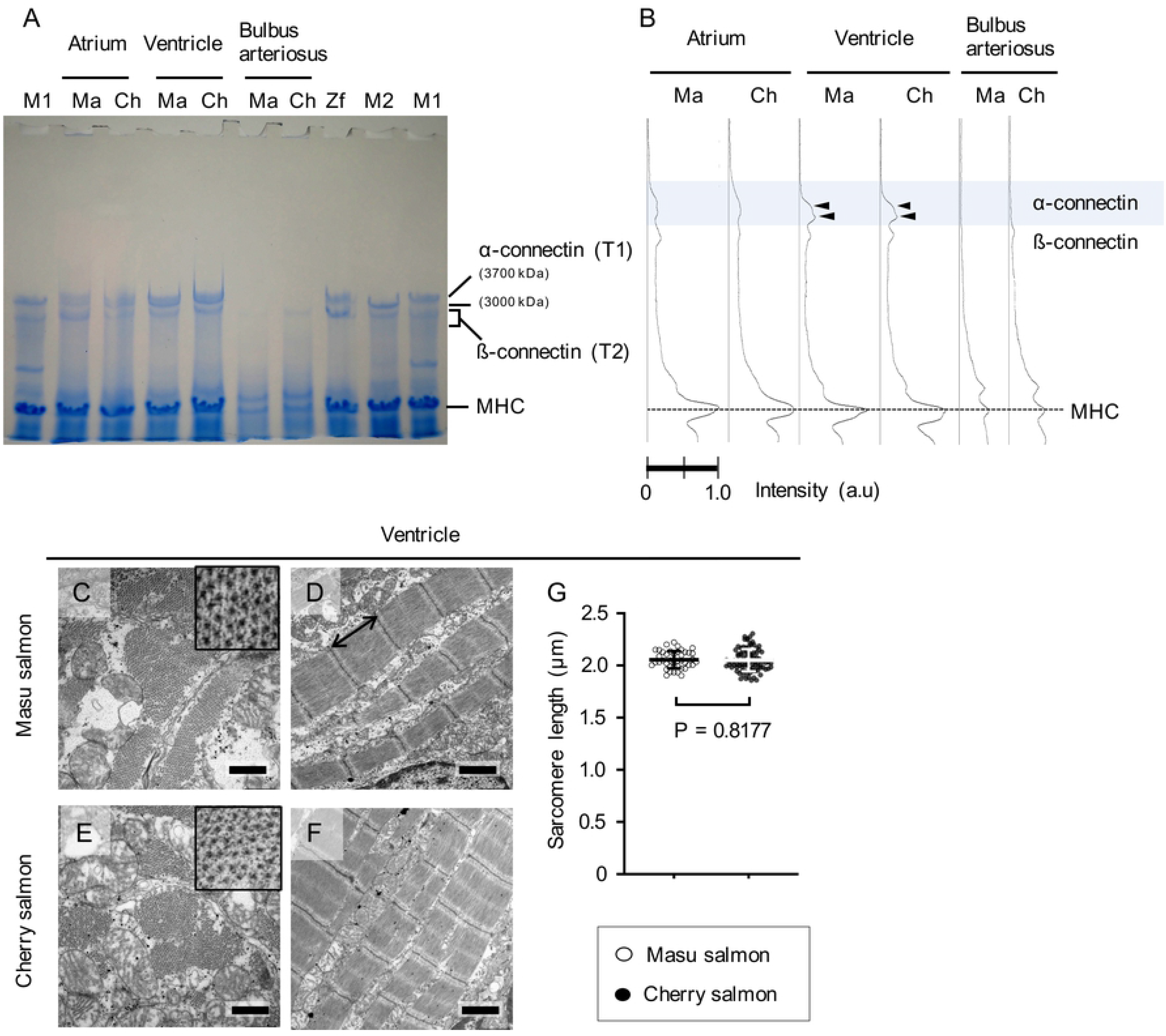
Molecular weights of the Connectin isoforms and sarcomere lengths in the hearts of masu and cherry salmon. (A) A representative Coomassie Brilliant Blue (CBB)-stained gel image. Lanes 2–7 show the molecular weights of the Connectin isoforms in the atrium, ventricle, and bulbus arteriosus of masu salmon (Ma) and cherry salmon (Ch). The following samples were used as molecular weight standards: a major Connectin isoform in zebrafish heart (Zf, 3500 kDa) in lane 8; Connectin N2A isoform from the skeletal muscles of mice (M1, 3700 kDa) in lanes 1 and 10; the Connectin N2B isoform expressed in the left ventricle of mice (M2, 3000 kDa) in lane 9. α-connectin (T1) and β-connectin (T2) indicate an intact Connectin and its degraded product, respectively. MHC indicates Myosin heavy chain (molecular weight ∼220 kDa each), which was used as a loading control. (B) Electropherogram of lanes 2–7 of the CBB-stained gel in (A). Black arrowheads indicate the band peaks for the ventricles of masu and cherry salmon in the α-connectin zone. (C–F) Representative transmission electron microscopy images of the cross-sections and longitudinal sections in the ventricular sarcomeres of masu and cherry salmon 29 months post fertilization. Insets of (C, E) showing higher magnification images of the cross-sections of myofibril bundles. Scale bars = 500 nm in (C) and (E) and 1 µm in (D) and (F). (G) The ventricular sarcomere lengths of the z-line to z-line in the ventricles of masu salmon (2.06 ± 0.08 µm, N = 42) and cherry salmon (2.05 ± 0.11 µm, N = 90). A black double arrow indicates sarcomere length. Lines and error bars indicate the means ± standard deviations. ○: masu salmon, ●: cherry salmon.

The TEM images of heart tissues from the masu and cherry salmon were observed and the length of the sarcomeres was measured (Fig 5C–G and S10A–E Fig). The sarcomere length of the ventricular myocardium did not vary significantly between the masu (2.06 ± 0.08 µm) and cherry salmon (2.05 ± 0.11 µm; *p* = 0.8177, Fig 5G). Similarly, the sarcomere length of the atrial myocardium also did not vary significantly between the masu (2.13 ± 0.16 µm) and cherry salmon (2.12 ± 0.11 µm; *p* = 0.7353, S7E Fig). In other words, the sarcomere lengths of the masu and cherry salmon under resting tension were similar. The bulbus arteriosus of the teleosts is mainly composed of abundant extracellular matrixes and smooth muscle cells [87–89]. Moreover, microfilaments and collagen fibers were observed in the bulbus arteriosus of both the masu and cherry salmon (S10F–H Fig) [90].

## Discussion

### Differing ventricular diastolic hemodynamics in masu and cherry salmon

In this study, pulsed-wave Doppler echocardiography revealed that landlocked masu salmon and sea-run cherry salmon have different ventricular diastolic properties despite being the same age and members of the same species. In masu salmon, monophasic atrioventricular inflow waveforms were observed immediately after the P wave in the electrocardiograph (Fig 1C), and therefore, these inflows may be attributed to atrial contraction. Simultaneous measurement of atrial and ventricular pressures showed that the atrial-ventricular pressure gradient occurred during atrial systole but not the early diastolic phase between completion of ventricular ejection and atrial systole (S4 Fig). These results suggested that there was no ventricular suction of blood from the atrium in masu salmon. Conversely, cherry salmon mainly displayed two types of atrioventricular inflow patterns, which were e/a fused and biphasic waveforms, depending on the heart rate (Fig 1D and 1E, S4 Fig). The inflows observed before detecting the P wave indicated passive ventricular filling, suggesting that the ventricle in cherry salmon acquired the ability to suction blood from the atrium. However, the atrial systolic velocity was greater than the early diastolic velocity (Fig 1F, E/A ratio = 0.48 ± 0.06); therefore, it was clear that atrial contraction plays a dominant role in ventricular filling in cherry salmon, at least under anesthesia.

A negative correlation has been reported between body mass and resting heart rate in both mammals and birds [91, 92], but this rule does not always apply to fish [93]. However, in our study, at 29–30 mpf, cherry salmon was over 10 times heavier than masu salmon (Table 1), and the heart rate of masu salmon was placed higher than that of cherry salmon (S1 Table, *p* = 0.0164). A previous hemodynamic study of rainbow trout had suggested that the heart rate is associated with the appearance of monophasic or biphasic atrioventricular inflow waveforms [94]. Pulsed-wave Doppler echocardiography of rainbow trout recorded biphasic waveforms in specimens with a low heart rate (44–69 bpm) [94]. In cherry salmon, monophasic atrioventricular inflow switched to biphasic when the R-R interval times extended, but the time duration between the end of ventricular ejection and the onset of new inflow was constant at 92−96 ms, independent of the R-R interval time (S4 Fig). In comparison, the lower heart rate cherry salmon had longer ventricular diastole, and the event of ventricular filling of blood occurred before atrial contraction began; however, no biphasic inflow waveform was observed in masu salmon, even though the time duration from ventricular ejection to the onset of atrioventricular inflow increased. However, since juvenile Atlantic salmon of 15 g body weight showed biphasic waveforms [95], it is possible that our data of pulsed-wave Doppler echocardiography are *O. masou*-specific. Elastic reaction forces, muscle contraction, and differences in chamber size between the atria and ventricles have been considered as candidates for the driving forces of ventricular filling, but they have not been sufficiently understood yet [96, 97]. Our study revealed that the same species of fish adapted to different environments, river and sea, acquired different ventricular filling systems during diastole. Thus, histological and material changes in the ventricles of cherry salmon might have contributed to generating the early diastolic ventricular inflow (Figs 2 and 3).

### Relative ventricular stiffness index greater in cherry salmon than masu salmon

The mechanical properties of ventricles of different sizes cannot simply be evaluated by comparing the individual mechanical parameters [4]. To assess ventricular contraction, end-systolic pressure–volume analysis had been carried out by normalizing ventricular volume per myocardial mass with respect to its contraction indexes [98]. EDPVR analysis, based on a time-varying elastance model, is an important method to assess ventricular stiffness [4]; however, its application in comparing the stiffness of ventricles of different sizes is difficult. In a smaller ventricle, the ratio of the pressure increase relative to the volume increase is greater than for a larger one. Thus, the EDPVR curve for a smaller heart would be steeper than that for a larger ventricle, even with the same actual stiffness. In this study, to evaluate the relative ventricular stiffness of masu and cherry salmon, we calculated the stiffness per unit of myocardial mass by normalizing the ventricular end-diastolic volume by the ventricular mass [15]. The index of ventricular stiffness per unit myocardial mass of cherry salmon, calculated from the normalized EDPVR curve (Fig 2A), was significantly higher than that of masu salmon (Fig 2B), thus suggesting that the ventricle of cherry salmon has a relatively suppressed diastolic function.

The ventricular walls of cherry salmon were thickened, and its coronary vascular network was well developed, as compared to those of masu salmon (Fig 3D and 3H). According to Laplace’s law, the thicker the ventricular wall, the greater the wall tension generated in response to increased internal pressure, and the greater the ventricular stiffness. Physiological hypertrophy in the ventricular wall, which causes both stroke volume and systolic pressure to increase, improved the overall cardiac output [99, 100]. Therefore, despite the expected high performance of cherry salmon hearts, their ventricles were stiffer than those of masu salmon (Fig 2). Coronary circulation presents hemodynamic characteristics that are not observed in other organs: during ventricular systole, arterial blood flow is blocked because of the pressure associated with contraction, while during diastole, blood flows only after the percutaneous pressure is reduced. Therefore, excessive stretching of the coronary vessels associated with ventricular dilation increases vascular resistance and decreases blood supply to myocardial tissues. Thus, the stiffening of the ventricle in cherry salmon might support the coronary vessel shapes and set the upper threshold for stroke volume to avoid excessive ventricular filling to supply sufficient amounts of arterial blood to the myocardium through the coronary network.

### Collagen accumulation in the thick compact layer of cherry salmon

In this study, histological analysis provided the causes of ventricular stiffness. The ventricle of cherry salmon had a thicker compact layer compared to that of masu salmon (Fig. 3D, 3H, 3K). This is consistent with previous reports on other salmonids wherein the compact layer thickens with growth [56, 57]. This histological change indicates that the growth ratio of the compact and sponge layers was different during the growth process of the masu and cherry salmon. There were no differences in the morphology and nuclear number of each isolated cardiomyocyte between the masu and cherry salmon (Fig 4; S7 and S8 Figs). Therefore *O. masou* hearts may show growth because of proliferation of cardiomyocytes or the differentiation of progenitor and stem cells instead of cardiomyocyte hypertrophy.

The compact layer of cherry salmon also contained more collagen fibers as compared with masu salmon (Fig 3E, 3I, and 3M). Increased stiffness in the ventricle correlates with tissue fibrosis [66, 101], and is reportedly induced by collagen accumulation in the myocardium [102]. In a rainbow trout study, breeding at lower temperatures was found to reduce ventricular compliance mediated by accumulating collagen fibers but not because of hypertrophy of the compact layer [65]. Hence, the stiffening of the ventricle in cherry salmon is assumed to be strongly correlated to increasing collagen fiber content in the compact layer.

### Similar expression patterns of Connectin isoforms in the ventricles of masu and cherry salmon

The expression ratios of the Connectin isoforms affect cell passive tension [30, 35–37]. The mammalian heart expresses the lower-molecular-weight Connectin isoform N2B and the higher-molecular-weight Connectin isoform N2BA [31–33]. Rainbow trout express N2B-like and N2BA-like proteins in their hearts [103]. Multiple *connectin* splicing variants are reportedly expressed in zebrafish [104]. The SDS-agarose gel electrophoresis in this study expected the expression of two types of Connectins in the ventricles of the masu and cherry salmon (Fig 5A and 5B); however, western blotting could not discriminate between these Connectins because either one of the isoforms might not have been detected by the antibody used in this study (S10 Fig). The similarity in the expression pattern of Connectins in the ventricles of cherry and masu salmon indicated that the difference in ventricular stiffness was not because of regulation by splicing of connectin molecules. There was no difference in sarcomere length under unloaded conditions as assessed by TEM (Fig 5C–G). However, the correlation between the molecular sizes of Connectins and sarcomere length is controversial [105, 106]. Although the regulation of tensile strength by Connectin phosphorylation is a possible mechanism other than the expression of its isoforms [27], we did not analyze it. In addition, we did not identify the gene and amino acid sequence of the Connectins. Salmon have undergone four genome duplication events during their speciation, and consequently, their genomic DNA sequences are extremely complex and difficult to analyze [107–109]; however, recently, the complete genome of the Atlantic salmon was sequenced [79]. *O. masou* is also assumed to have a complex genome, but its genomic DNA sequences have been gradually deciphered, and comprehensive gene expression analysis is being attempted [110].

In our experiment, we used *O. masou* of the same age but did not consider sex because it could not be specified at the time of purchase; however, previously, differences between male and female wild-type zebrafish strains at 9 mpf were reported when considering heart morphology, heart rate, and cardiac function parameters [111]; furthermore, rainbow trout have been found to differ based on sex in the amount of connective tissues in their ventricles [112]. We used non-strained *O. masou* with non-uniform genetic backgrounds in this study, but a rigorous experimental system with a molecular basis is needed to provide definite answers to questions regarding the processes of heart development, morphogenesis, functions, and gene expression patterns.

## Conclusion

In this study, the properties of ventricular diastole and heart histology were analyzed in landlocked masu and sea-run cherry salmon at 29–30 mpf. Because cherry salmon had thick ventricular walls and accumulated collagen in the myocardium, their ventricles were found to be stiffer than those of masu salmon on a per unit myocardial mass basis. Histological adaptations of the heart because of body growth and the living environment cause hemodynamic changes in *O. masou*, and the atrioventricular inflows of cherry salmon showed biphasic waveforms with the early diastolic filling and the atrial contraction. Our study is expected to provide evidence of the vertebrate ventricle acquiring its suction function by modulating the stiffness of the myocardium and changing the ventricular filling system during development, environmental change, speciation, and evolution.

## Acknowledgments

The contributing authors would like to thank Nobuhisa Iwachido (Kawasaki Medical School, Japan) for their technical assistance with the tissue staining and Nobuaki Matsuda (Kawasaki Medical School, Japan) for operating the electron microscope. The authors would like to thank Editage for English language editing.

## Supporting information

**S1 Fig. Anatomical observations of a fish heart**

(A) An image of the left lateral view of a masu salmon heart at 29 months post fertilization (mpf). Scale bar = 1 cm. (B) Magnified view of the white box in panel (A). At, atrium (green dashed line); V, ventricle (magenta dashed line); Ba, bulbus arteriosus (yellow dashed line); Sv, sinus venosus (white dashed line); Di, diaphragms (blue dashed line); Li, liver. (C) Life history of *Oncorhynchus masou masou*. (D) A juvenile *O. masou* at 6 mpf. Scale bar = 1 cm. (E) Stereomicroscopic images of a juvenile heart. Scale bar = 1 mm. (TIFF)

**S2 Fig. Simultaneous echocardiographic and electrocardiographic measurements of Oncorhynchus masou masou**

(A) Method used for the fish echocardiography and electrocardiography. To record the cardiac dynamics on the sagittal axis, anesthetized fish were turned upside down and secured to the holder, and the transducer probe was vertically and directly positioned above the heart. Electrodes were clipped to the pectoral fins and pelvic fin. Air was supplied continuously during the experiments. (B, C) Echocardiographic images of the sagittal axis; (B) masu salmon at 30 months post fertilization (mpf); (C) cherry salmon at 30 mpf. At, atrium (green dashed line); V, ventricle (magenta dashed line); Ba, bulbus arteriosus (yellow dashed line); Li, liver. (D) Electrocardiography results from the body surface of the masu salmon. P: P wave, QRS: QRS complex, T: T wave.

(TIFF)

**S1 Movie. Longitudinal axis echocardiography of a masu salmon**

(mp4)

**S2 Movie. Longitudinal axis echocardiography of a cherry salmon**

(mp4)

**S1 Table. Heart rates of the masu and cherry salmon**

(docx)

**S3 Fig. Atrioventricular inflow waveforms at different R-R interval times**

(A) Pulsed-wave Doppler images of masu (left column) and cherry salmon (right column). Upper row; representative ventricular ejection waveform (326.75 ± 1.04 ms in masu salmon #5 and 351.75 ± 1.09 ms in cherry salmon #1). Second−lower rows; atrioventricular inflow waveforms observed at short, intermediate, and long R-R interval times. The yellow line indicated the average time from the peak of the R wave to the end of ventricular ejection. The yellow double-headed arrows indicated the time from the end of ventricular ejection to the onset of atrioventricular inflow. (B) Times of R-R interval and from the end of ventricular ejection until atrioventricular inflow were observed in each panel in (A).

(TIFF)

**S4 Fig. *In vivo* atrial-ventricular pressure analysis of masu salmon**

(A, B) Recordings of the ventricular pressure in the masu salmon at 34 mpf. Ventricular and atrial pressures during three heart cycles (A). Ventricular and atrial pressure in the ventricular diastole (B); magnification of the 1 and 2 s range of panel (A). The solid and dotted lines indicate the ventricular pressure and atrial pressure, respectively. VD, ventricular diastole; IR, isovolumic relaxation; AS, atrial systole.

(TIFF)

**S5 Fig. End-diastolic pressure–volume relationship analysis for masu and cherry salmon**

(A) The horizontal axis shows the ventricular chamber volume normalized by the mass of the masu salmon ventricles (N = 4) and the cherry salmon ventricles (N = 3). (B) Ventricular mass and index of ventricular stiffness in the masu and cherry salmon. The results of the EDPVR (A) were applied to Equation (1) to obtain the exponential *C*, ○: masu salmon, ●: cherry salmon.

(TIFF)

**S6 Fig. Hematoxylin and eosin staining of isolated fish hearts**

(A–H) Hematoxylin and eosin staining images of the sagittal sections of the hearts at 29 months post fertilization: masu salmon (A–D) and cherry salmon (E–H). (A, E) Images of the whole heart. Scale bars = 5 mm. (B, F) Magnified images of the ventricles. Scale bars = 1 mm. (C, G) High-magnification images of the compact layers. Scale bars = 100 µm. (D, H) High-magnification images of the spongy layers. Scale bars = 100 µm. The cytoplasm is shown in magenta, and the nuclei are shown in blue–purple. At, atrium; V, ventricle; Ba, bulbus arteriosus; Cl, compact layer; Sl, spongy layer.

(TIFF)

**S7 Fig. Immunofluorescence staining of cardiomyocytes**

(A–F) Immunofluorescence images show attached cardiomyocytes from the masu and cherry salmon at 29 months post fertilization. Cardiomyocytes were stained with antibodies recognizing Alpha-actinin-2 at z-lines (green) and Hoechst 33342 to detect nuclei (blue). (B, D) Magnifications of the images in (A) and (C). (E, F) Merged bright-field and immunostaining images. (F) Non- cardiomyocyte; magnification view of the white dotted line enclosure in (E). Scale bars = 20 µm in (A), (C) and (E), and 5 µm in (B), (D) and (F).

(TIFF)

**S8 Fig. A binucleated cardiomyocyte**

(A) A Hoechst 33342-stained binucleated cardiomyocyte (blue) in a cherry salmon. (B) Magnified image shows Hoechst 33342 signals in the grayscale mode. Scale bars = 20 µm. (TIFF)

**S9 Fig. Anti-Connectin antibody immunoblots**

Representative and original immunoblot image showing the detection of Connectin expression in masu (Ma) and cherry (Ch) salmon hearts. Connectin N2A isoform in the mouse skeletal muscle in lanes 1 and 9 (M1, 3,700 kDa) and the N2B isoform in the left ventricle of mice in lanes 2 and 10 (M2, 3,000 kDa) were used as a positive control for the experiment and as a guide for molecular weight. α-Connectin (T1) and ß-connectin (T2) indicate intact Connectins and their degraded products, respectively.

(TIFF)

**S10 Fig. Electron micrographs of masu and cherry salmon hearts**

(A–D) Representative transmission electron microscopy images of the cross-sections and longitudinal sections of the atrial sarcomeres of masu and cherry salmon at 29 months post fertilization (mpf). Insets of (A, C) showed higher magnifications of cross-sections of the myofibril bundles. (E) The graph showed atrium sarcomere lengths of the z-line to z-line in masu salmon (2.13 ± 0.16 µm, N = 25) and cherry salmon (2.12 ± 0.11 µm, N = 103). One or two ultrathin slices were observed in each tissue sample. Lines and error bars indicate means ± standard deviations. ○: masu salmon, ●: cherry salmon. (F–H) Representative transmission electron microscopy images of the bulbus arteriosus of masu and cherry salmon at 29 mpf. (G) Higher magnification image in the black dashed line in (F). FM: microfilament; CF: collagen fiber Scale bars = 500 nm in (A, C), 1 µm in (B, D), and 2 µm in (F–H).

(TIFF)

